# Hidden regenerative state in planarians: A geometric model of bioelectric memory using Tangential Action Spaces

**DOI:** 10.64898/2026.04.01.715890

**Authors:** Marcel Blattner

## Abstract

Planarian fragments can regenerate with normal gross anatomy after a transient bioelectric perturbation yet display altered outcomes upon re-cutting, implying that regeneration can store a persistent hidden state. Here we formulate an open-path version of Tangential Action Spaces (TAS) for this setting. Regeneration after a given cut is represented as a prescribed coarse anatomical trajectory together with multiple physiological lifts in a higher-dimensional state space. A metric on physiological state space defines a baseline lift, an effective excess-cost functional, and a baseline-relative endpoint displacement that serves as written hidden regenerative state.

Re-cutting converts this open-path construction into a challenge readout. Locally, the theory yields a cut-dependent memory co-metric that identifies latent directions that are easy, difficult, or inaccessible to rewrite. We show that this geometry is consistent with published observations of cryptic phenotypes, stable re-challenge ratios, and near-absorbing double-headed outcomes. A reduced rank-one latent-threshold realization fitted to published 8-OH immediate and re-challenge counts identifies a challenge-sensitive cryptic interval below the immediate double-headed threshold and predicts out-of-sample re-challenge penetrances near 15% for nigericin- and monensin-treated immediate single-headed survivors using only their immediate phenotype penetrances.

As a mechanistic bridge, a local electrodiffusive in-silico example instantiates a local version of the physiological-state effort metric *G*. This metric defines the baseline lift and excess rewriting cost, in relative biophysical units, and yields explicit example local write geometry. An illustrative semimechanistic readout based on integrated wound-edge gap-junction contrast and Na/K-ATPase load reproduces the treated-family ordering and similar transfer predictions when the untreated baseline is softly anchored near zero. These quantitative layers are intended as proof-of-concept calibratability and mechanistic-grounding checks rather than full validation of the complete open-path model. The framework therefore turns cryptic regenerative memory into a geometric, costed, and experimentally testable object, yielding predictions about temporal-profile dependence, compensatory cancellation, sign-reversing controls, cut dependence, anisotropic rewriting, and multi-round accumulation of hidden regenerative state.

## 1 Introduction

Planarian regeneration is robust but not fixed. Following amputation, body fragments reliably rebuild missing structures, yet transient perturbations of bioelectric state during the earliest phases of regeneration can redirect the long-term patterning outcome [1–7]. In some cases the effect is immediately visible, producing stable double-headed (DH) animals. In other cases the fragment regenerates as an apparently normal single-headed (SH) worm, but a later standardized re-cut reveals an abnormal SH/DH outcome distribution. This second case implies that the first regeneration round has written a persistent hidden state that is not resolved by gross anatomy alone.

That phenomenon has been described as a *cryptic phenotype* [4]. Cryptic worms are histologically similar to wild type and can show normal expression of key polarity markers and normal neoblast distributions, yet they retain a non-genetic memory of prior perturbation that becomes visible only under challenge [4]. Existing experimental work points to a distributed pattern-control system involving bioelectric state, gap-junctional coupling, and muscle-resident positional information [2, 8–12]. What is still missing is a quantitative language that cleanly separates three objects: the visible anatomical trajectory of regeneration, the hidden physiological state retained after regeneration, and the re-cut assay that reveals that hidden state.

Here we adapt Tangential Action Spaces (TAS) [13] to provide that language. TAS is a geometric framework for systems that follow a prescribed coarse trajectory while retaining degrees of freedom that are invisible at the coarse level. For planarian regeneration, the coarse object is a cut-specific regeneration trajectory in a low-dimensional anatomical state space; the hidden object is the endpoint displacement within a higher-dimensional physiological fibre over the same visible endpoint. In this setting, memory means a persistent displacement of the latent regenerative controller relative to a baseline regenerative episode, even when the first-round anatomy appears normal.

This perspective is useful for three reasons. First, it gives a formal definition of hidden regenerative state. Second, it separates the baseline cost of executing a regeneration trajectory from the additional effective intervention cost required to write a hidden shift. Third, it yields a cut-dependent local geometry of writable and non-writable directions in latent state space, which can be probed experimentally by standardized challenge cuts.

The manuscript has three aims. The first is theoretical: to formulate an open-path version of TAS appropriate for regeneration, where the relevant object is a regeneration episode from a post-cut state to a homeostatic anatomy rather than a closed mechanical loop. The second is empirical: to show that a reduced latent-threshold version of the model can already be constrained by published phenotype counts. Using published 8-OH immediate and challenge data as a calibration set, we estimate a lower challenge threshold beneath the immediate DH threshold and then generate out-of-sample predictions for the re-cut penetrance of nigericin- and monensin-treated immediate SH survivors using only their immediate phenotype penetrance. The third is mechanistic and illustrative: to show a concrete local route by which *G* can be instantiated within an electrodiffusive model and coupled back to the same latent phenotype layer. In that third layer we use a BETSE-based local electrodiffusive example [14] to instantiate *G* in relative biophysical units and to test whether a mechanistically interpretable simulator-derived feature subset can recover the same treated-family ordering and transfer scale as the reduced free-*x*_k_ fit.

The goal is not to replace mechanistic bioelectric models of regeneration [12, 14–17]. Rather, it is to add a complementary geometric layer that defines hidden state, effective writing cost, and challenge readout in a common framework. Accordingly, the quantitative results below should be read in two empirical layers: a proof-of-concept reanalysis of published phenotype counts, and a local in-silico grounding bridge. Neither layer is a definitive parameter-estimation study of the full regenerative controller. Because currently available experimental data are compressed largely to endpoint phenotype counts, the quantitative sections below test a compressed rank-one projection of the full TAS geometry rather than the full open-path model. The more specifically geometric content of TAS, including cut dependence, anisotropic rewriting, and compensatory cancellation, therefore enters most directly through the higher-rank predictions and the local mechanistic bridge.

**Figure 1:**
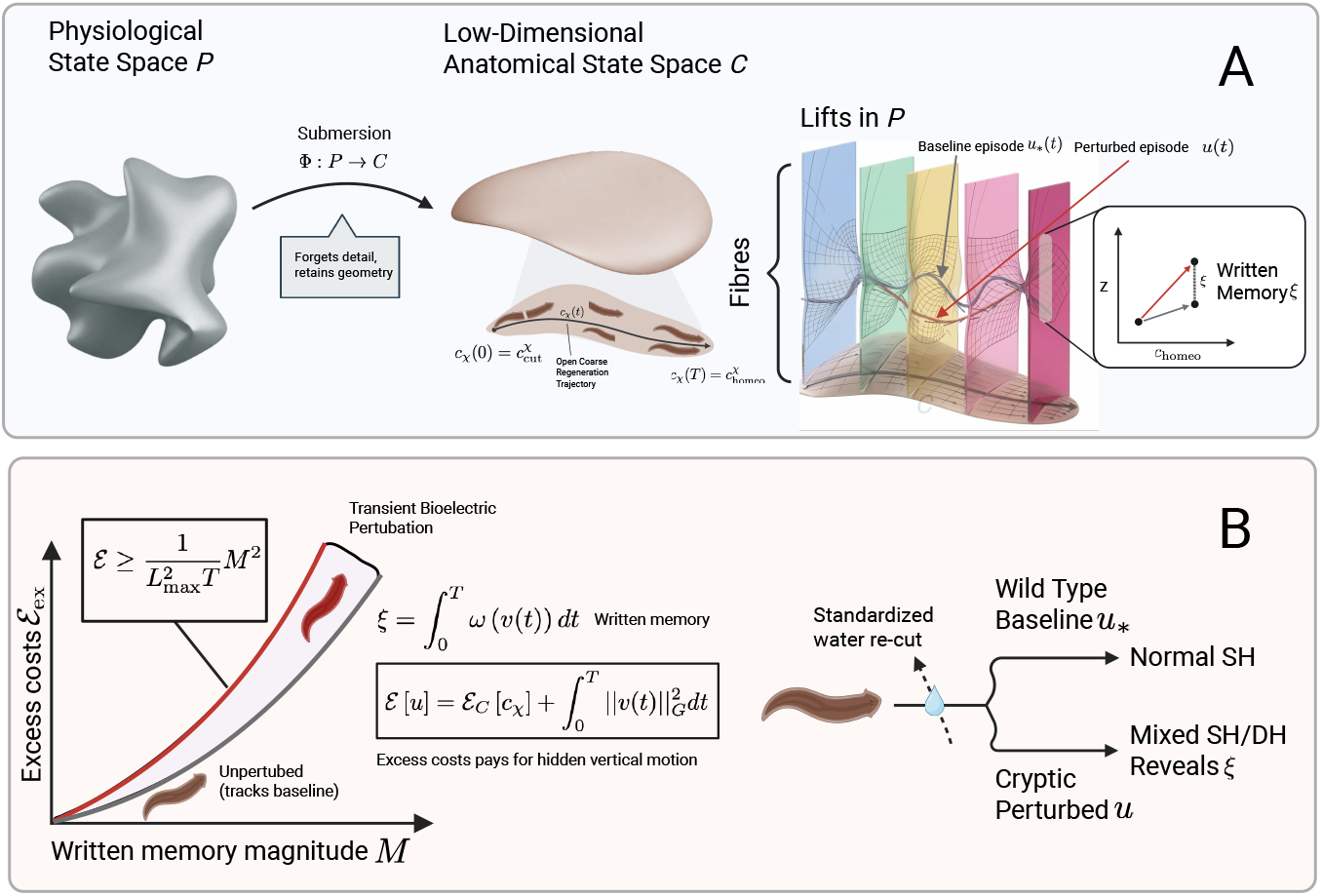
Overview of hidden regenerative state, effective writing cost, and challenge readout. **(A)** A regenerating fragment follows a prescribed coarse regeneration trajectory *c*_χ_(*t*) in a low-dimensional anatomical state space *C*, obtained from the full physiological state space *P* by a submersion Φ : *P* → *C*. Multiple physiological lifts of the same coarse trajectory exist in *P*. The physiological-state metric *G* selects a baseline lift *u*_∗_(*t*), while a transient bioelectric perturbation can generate a perturbed lift *u*(*t*) that reaches the same visible endpoint 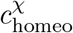 but ends at a displaced point on the final fibre. This baseline-relative endpoint displacement is the written hidden regenerative state *ξ*. **(B)** In the open-path TAS formulation, excess cost above baseline pays only for hidden vertical motion and is bounded below quadratically in the memory magnitude *M* = ∥*ξ* ∥_2_, yielding a local quadratic cost law. Biologically, this distinguishes wild-type baseline regeneration from a cryptic perturbed state: both can appear immediately normal single-headed, but a standardized water re-cut reveals the hidden shift by producing a mixed single-headed/double-headed outcome distribution from the cryptic branch.

## 2 A geometric framework for hidden regenerative state

### 2.1 State space and coarse regeneration trajectory

Let *P* denote the space of full physiological states of the regenerating animal and let *C* denote a manifold of coarse regeneration states. We assume a smooth surjective submersion

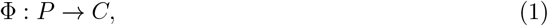

which forgets fine-grained physiological detail while retaining the coarse anatomical time course of interest. A point *p* ∈ *P* may be written locally as

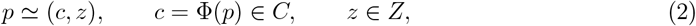

where *z* parametrizes latent pattern-control variables. For a first model we take *Z* ≃ ℝ^r^, acknowledging that a field-valued extension may ultimately be needed.

For a fixed cut type *χ* and regeneration horizon *T >* 0, let

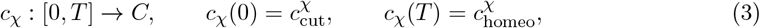

be a prescribed coarse regeneration trajectory from the standard post-cut state to the target homeostatic anatomy. A *physical regeneration episode* is any lift

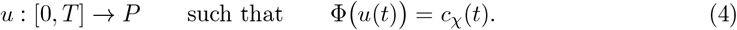

This is the regenerative analogue of a task trajectory in TAS. What re-cutting later exposes is not a failure to reach *c*_χ_(*T*), but the fact that different lifts of the same coarse trajectory can end at different hidden states on the fibre over 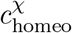.

Let

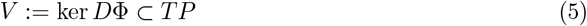

be the vertical bundle of task-invisible directions. We equip *P* with a Riemannian metric *G* that prices regenerative effort or effective intervention effort, and choose the metric connection

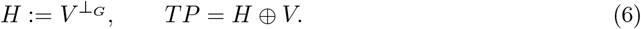

*Remark* 2.1 (Choosing the coarse regeneration state). The coordinates of *C* need not be limited to final anatomy. A useful coarse regeneration trajectory can include quantities such as wound closure, blastema growth, polarity-marker dynamics, organ-position markers and gross anatomical restoration [1, 5, 18–20]. The framework only requires that these variables define a reproducible low-dimensional path *c*_χ_(*t*) whose different physical lifts can be compared.

### 2.2 Metric lift, baseline episode and excess cost

At each *p* ∈ *P*, the metric lift is the minimum-*G*-norm velocity compatible with a given coarse velocity 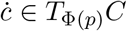:

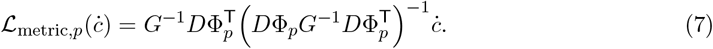

This induces a coarse metric on *C*,

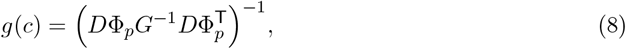

written locally on the operating domain where the fibre dependence is negligible.

For a chosen initial fragment state 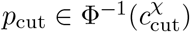, the *baseline regenerative lift u*_⋆_ is the metric lift of *c*_χ_ with *u*_⋆_(0) = *p*_cut_. Any other lift *u* with the same initial condition decomposes as

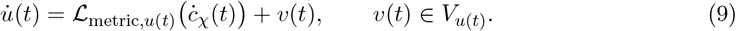

The total cost of an episode is

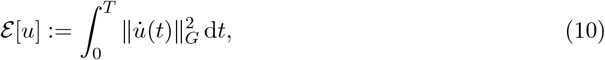

and the baseline coarse cost is

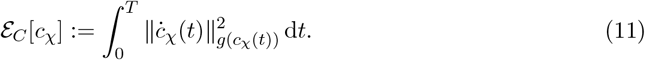

**Result 2.2** (Open-path baseline/excess decomposition). *For any lift u of c*_χ_,

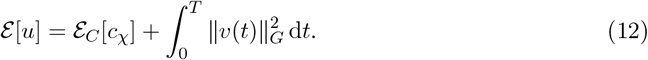

*The excess cost above baseline is therefore*

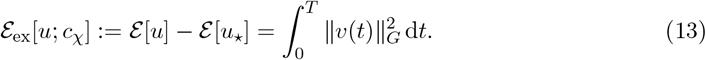

This is the regenerative analogue of the TAS accounting identity: baseline cost pays for realising the prescribed coarse regeneration trajectory, while excess cost pays only for hidden motion within the fibre. When unperturbed regeneration already tracks the baseline episode, all experimentally imposed effective effort appears in ℰ_ex_.

Here ℰ_C_[*c*_χ_] should not be read as the worm’s literal total metabolic cost of regeneration. It is the minimum of the chosen effective effort functional over all physiological lifts that realise the same coarse regeneration trajectory. Accordingly, ℰ_ex_ measures perturbation-induced hidden rewriting relative to that model-relative baseline, not total organismal energy expenditure.

### 2.3 Written regenerative memory for one episode

To define written memory, assume the final fibre over 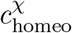 is represented locally by an Abelian coordinate ℝ^r^ and let

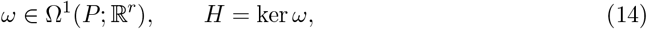

be the corresponding local connection one-form. The endpoint displacement of a lift relative to the baseline is then

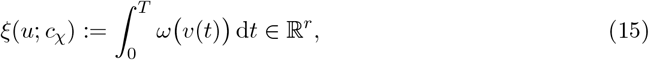

with magnitude

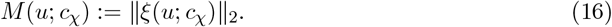

This is the open-path analogue of baseline-relative holonomy: the prescribed coarse trajectory is fixed, but the endpoint on the final fibre may shift.

**Result 2.3** (Open-path regenerative TAS law). *Let*

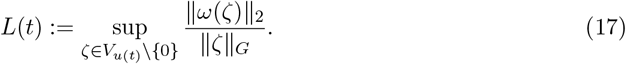

*Then every regenerative episode satisfies*

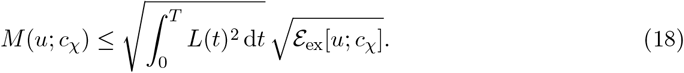

*If L*(*t*) ≤ *L*_max_ *on* [0, *T*], *then*

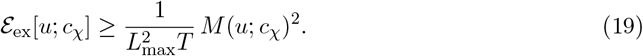

The proof is the same as in the closed-loop TAS setting: it uses only the path length of the relative endpoint motion on the final fibre and therefore does not require the base trajectory itself to be closed.

### 2.4 From open regenerative episodes to return maps

Re-cutting turns the open-path construction above into a cycle-to-cycle map. Let

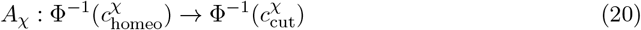

be the cut operator for injury type *χ*. For a pre-cut state *p* and a perturbation protocol labelled by *η*, let 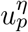 denote the lift of the same coarse trajectory *c*_χ_ starting from *A*_χ_(*p*). The one-cycle map is

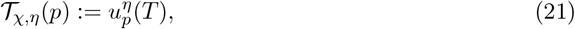

with baseline map 𝒯_χ,⋆_ := 𝒯_χ,0_. Thus the hidden state written in a single regenerative episode is the relative fibre displacement between 𝒯_χ,η_(*p*) and 𝒯_χ,⋆_(*p*) over the same visible endpoint 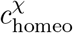.

Let Π : *P* → *Y*_obs_ be a phenotype readout. The immediate readout is *R*_imm_ := Π. Given a standardized challenge cut *χ*_ch_, define the challenge readout

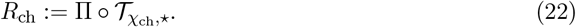

Linearizing at the baseline endpoint yields

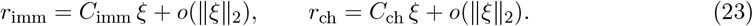

A hidden shift *ξ*≠ 0 is therefore *cryptic* when

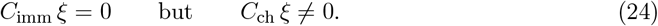

In words: the first round looks normal at the chosen resolution, but a standardized re-cut reveals that the latent regenerative controller has been rewritten.

### 2.5 Local write map and the induced memory co-metric

Fix a cut type *χ* and a *k*-parameter family of small perturbations *η* ∈ ℝ^k^ around the baseline regenerative episode. Assume the hidden vertical deviation linearizes as

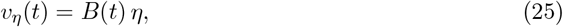

for some chosen vertical modes *B*(*t*) along the baseline lift. Then

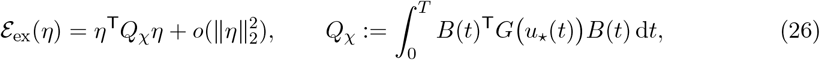

and the written regenerative memory linearizes as

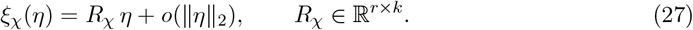

#### Definition 2.4

(Cut-dependent memory co-metric). *The induced memory co-metric is*

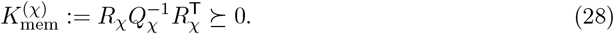

**Result 2.5** (Local regenerative write law). *Assume Q*_χ_ ≻ 0. *For any target hidden shift ξ* ∈ range(*R*_χ_), *the minimal excess cost required to write it in one regenerative episode is*

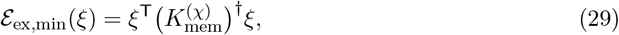

*where* (·)^†^ *denotes the Moore-Penrose pseudoinverse. Consequently*,

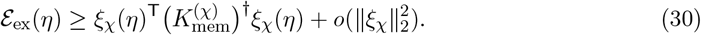

Large eigenvalues of 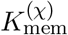 correspond to hidden directions that are easy to rewrite during regeneration from cut *χ*; small eigenvalues correspond to difficult directions, and the kernel identifies perturbations that dissipate cost with little or no first-order rewriting. Different cut types induce different 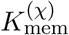 because the accessible latent directions depend on the geometry of the injury and the baseline regenerative episode.

The challenge readout inherits its own observable co-metric,

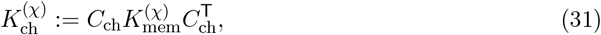

so that locally

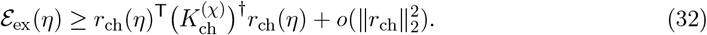

This is the directly testable version of the theory: cryptic memory need not be reconstructed internally if a standardized challenge readout resolves it.

### 2.6 Multi-round accumulation

If the latent controller is approximately retained from one episode to the next, written shifts accumulate across regeneration cycles. In the linear Abelian regime,

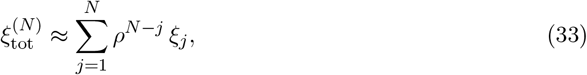

where *ρ* ∈ (0, 1] is a retention factor and *ξ*_j_ is the hidden shift written in round *j*. Thus repeated sub-threshold perturbations can push the latent state across a challenge threshold even when no single round does so on its own. If the latent controller is genuinely nonlinear or manifold-valued, this additive law is replaced by a transported composition rule analogous to the non-Abelian extension of TAS [13], so the order of perturbations becomes experimentally relevant.

## 3 Consistency with published bioelectric regeneration experiments

We now show how the open-path TAS construction is consistent with several characteristic findings from published planarian bioelectric regeneration experiments. The point is not that the framework predicts the exact numbers from first principles, but that it identifies the geometric structure those findings imply.

### 3.1 The cryptic phenotype

#### Finding (Durant et al. 2017)

Brief gap-junction blockade during regeneration of pharyngeal fragments produces a mixed population: some animals become double-headed (DH), while others regenerate as apparently normal single-headed (SH) worms. When the SH worms are re-cut in plain water, they again produce a characteristic SH/DH mixture [4].

#### TAS interpretation

The first regenerative episode follows the same coarse anatomical path *c*_χ_(*t*) to the same visible endpoint, but some lifts end at a hidden displacement *ξ* ≠ 0 on the final fibre. For the SH fraction, this displacement lies in ker *C*_imm_ and is therefore invisible to the immediate readout. Upon re-challenge, however, *ξ* ∉ ker *C*_ch_, so the hidden shift is converted into a visible change in regenerative outcome. What experiments call a cryptic phenotype is therefore precisely a challenge-visible but immediately invisible direction in the latent regenerative controller.

### 3.2 Stable challenge ratios reflect latent state, not treatment heterogeneity

#### Finding

The characteristic SH/DH ratio after re-challenge is not explained by incomplete exposure of the original cohort. Rather, all animals were treated, and the altered ratio reflects a shared hidden change in regenerative state [4].

#### TAS interpretation

A treatment writes a similar hidden shift *ξ* in each worm, moving the cohort from the baseline endpoint into a region of latent space where the challenge map *R*_ch_ yields a mixed outcome distribution. The ratio is therefore a property of the challenge kernel evaluated at the shifted latent state, not a statement that only a subset of worms “received” the perturbation. In the scalar model below, this corresponds to writing the system into a cryptic interval that is immediate-readout silent but challenge-readout active.

### 3.3 Near-absorbing double-headed outcomes

#### Finding

Double-headed worms tend to regenerate as double-headed upon re-cutting, whereas cryptic single-headed worms can generate either wild-type or double-headed outcomes [4].

#### TAS interpretation

In a nonlinear latent-state picture, the DH phenotype corresponds to a second attractor basin of the regenerative controller. Cryptic worms sit near the basin boundary: they end one coarse regeneration episode at a hidden displacement that is not yet immediately visible, but the next challenge episode resolves that displacement stochastically. Once the latent state crosses deeply into the DH basin, subsequent baseline regenerative episodes simply reproduce that attractor. Thus the near-absorbing behavior of the DH state over the observed range reflects the nonlinear geometry of the latent controller, while the cryptic state reflects hidden displacement near a separatrix.

### 3.4 Different molecular perturbations can share an effective write direction

#### Finding

Distinct perturbations that depolarise wound tissue can generate similar cryptic/DH outcomes even though their molecular mechanisms differ [3–5].

#### TAS interpretation

Different perturbation modalities correspond to different families of lifts of the same coarse regeneration trajectory. If their first-order effect projects onto the same dominant write direction in the local map *R*_χ_, they can generate similar hidden shifts *ξ* despite acting through different biochemical pathways. The common outcome is therefore not “mechanism independence” in a literal sense; it is shared projection onto an effective regenerative write direction. This is the biological analogue of distinct nullspace controls writing the same memory in a redundant mechanical system when they share the same component along a common dominant latent channel.

### 3.5 Brief perturbations can permanently rewrite the controller

#### Finding

A brief perturbation during the first hours after amputation can suffice to induce long-lived cryptic or DH outcomes, even after complete washout of the treatment [2, 4–6].

#### TAS interpretation

The open-path formulation makes it natural that the timing of a perturbation matters through its alignment with the baseline regenerative episode *c*_χ_(*t*). If the local write gain is high during an early sensitive segment of that trajectory, then a short pulse delivered in that window can write the same hidden displacement as a longer, weaker protocol delivered elsewhere. Once the hidden shift has been written, the later autonomous portion of the regenerative episode transports it to the final fibre, where it persists until revealed by re-challenge.

## 4 A reduced scalar model

To make the framework concrete, consider the rank-one regime in which the dominant hidden write channel is effectively scalar. Let *θ*_n_ ∈ ℝ denote a latent anterior-posterior pattern variable at the start of regeneration episode *n*. Let *s*_n_(*t*) be the signed write signal along the dominant local channel during that episode. Then a simple stochastic update is

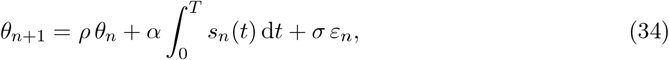

where *ρ* ∈ (0, 1] is retention, *α* is the write gain, *σ >* 0 is the cycle noise scale and *ε*_n_ ∼ 𝒩 (0, 1). The excess cost of the episode is

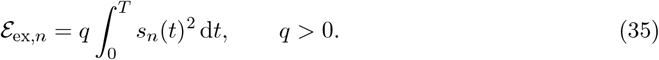

The hidden shift written in that episode is

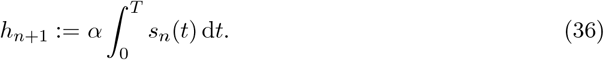

By Cauchy-Schwarz,

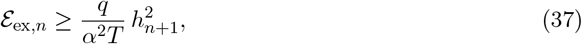

with equality when *s*_n_(*t*) is constant in time. This is the exact rank-one version of the open-path TAS law.

The immediate readout is taken to be thresholded anatomy,

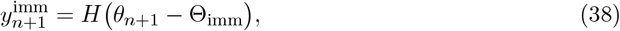

where *H* is the Heaviside step function. The challenge readout is more sensitive,

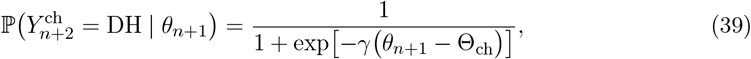

with Θ_ch_ *<* Θ_imm_. Hence the interval

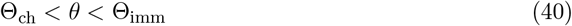

is cryptic: the worm looks normal after the first episode but has altered challenge probability. A nonlinear extension replaces the linear restoration term by a double-well potential,

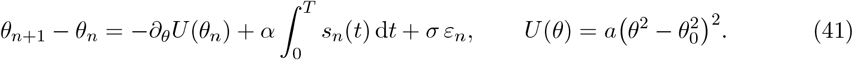

The two wells correspond to wild-type and double-headed attractors, while the cryptic state lies near the basin boundary. This minimal model is sufficient to represent hidden writing, immediate-readout silence, challenge-readout sensitivity and the near-absorbing behavior of the DH state over the observed range.

## 5 Calibration to published phenotype counts, an illustrative grounded in-silico bridge, and transfer predictions

The framework becomes empirically useful only if its parameters can be estimated or at least partially constrained. Here we pursue two deliberately limited empirical layers. The first is a reduced rank-one statistical realization fitted to published phenotype counts. The second is an illustrative local electrodiffusive bridge that instantiates *G* in relative biophysical units and supplies a simulator-derived feature map *h*_k_. The first layer tests proof-of-concept calibratability of the shared hidden-state / dual-readout structure; the second asks whether a mechanistically interpretable feature subset can recover the same treated-family ordering and transfer scale without free per-condition write effects. Neither layer by itself validates the full open-path geometry.

Because available experimental data are compressed largely to endpoint phenotype counts, the first empirical layer tests a compressed statistical shadow of the full TAS geometry rather than the full open-path model. We use the 8-OH re-challenge dataset of [4] as the primary calibration set because it contains both immediate and challenge outcomes. We then ask a sharper transfer question: after constraining the immediate and challenge thresholds on that dataset, can other depolarising interventions documented in [5] be placed on the same latent write axis using only their immediate DH penetrances, thereby generating quantitative predictions for re-cut outcomes not included in the calibration? In parallel, can a grounded local simulator-derived feature map recover the same treated-family ordering and similar transfer predictions without reintroducing free per-condition write effects? In the staged scheme below, the nigericin and monensin re-cut penetrances remain out-of-sample predictions.

### 5.1 Effective write coordinate and observation model

For a protocol condition *k* applied to cut type *χ*, let *u*_k_(*t*) denote the signed perturbation profile over the regenerative episode. In the rank-one approximation, its first-order projected write component is

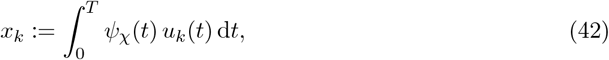

where *ψ*_χ_(*t*) is an unknown sensitivity kernel along the baseline coarse regeneration trajectory. If only endpoint data are available, one can replace Equation (42) by a free condition-level effect *x*_k_ or by a low-dimensional parametrisation tied to exposure duration and amplitude. The count-based reduced layer below uses this free-*x*_k_ route.

When early-window physiological measurements are available, one can go one step further and replace the abstract condition-level effect by a measured feature representation

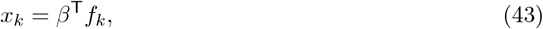

where *f*_k_ ∈ ℝ^m^ collects early-window bioelectric features and protocol covariates. A corresponding quadratic observable proxy for effective excess intervention cost is

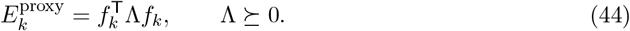

This preserves the reduced TAS geometry while making the dominant write coordinate and its associated cost partially observable from data.

A complementary route is mechanistic rather than purely statistical. Let *h*_k_ denote a vector of simulator-derived endpoint or early-window summaries produced by a local electrodiffusive model grounded in membrane voltage, transmembrane current, gap-junction flux and pump activity. One can then replace the free condition-level effect at the immediate layer by

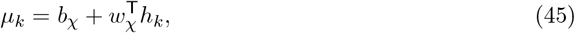

thereby coupling a grounded local *G* to the same latent readout layer. We use that strategy in Section 5.4 not as independent validation of the reduced fit, but as a semimechanistic compatibility check.

Let 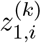 denote the latent regenerative controller of worm *i* after the first regeneration round under condition *k*. We model

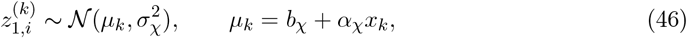

where *b*_χ_ is the cut-specific baseline offset, *α*_χ_ is the write gain and *σ*_χ_ collects between-worm heterogeneity and unresolved physiological fluctuations. Let *ϕ*_N_ and Φ_N_ denote the standard normal density and distribution function.

We take the immediate double-headed (DH) phenotype to occur when the latent controller crosses a threshold *θ*_imm_. Hence the immediate DH probability in condition *k* is

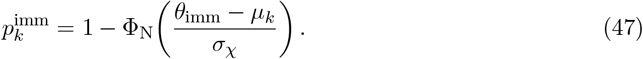

If *N*_k_ worms are scored after round 1 and *Y* ^imm^ are DH, then

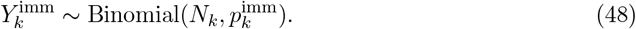

For the challenge readout, we model a baseline re-cut as propagating the latent state with retention factor *ρ*_χ_ ∈ (0, 1] plus challenge noise:

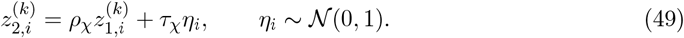

A challenged worm is scored DH if 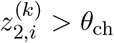, where typically *θ*_ch_ *< θ*_imm_. For worms that were single-headed (SH) after round 1, the conditional challenge DH probability is therefore

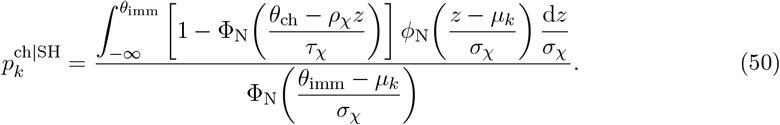

Similarly, for worms that were DH after round 1,

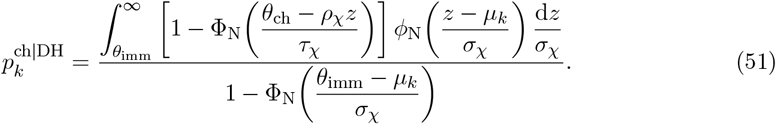

The second quantity is the formal version of the near-absorbing DH outcome check: over the fitted operating range, one should find 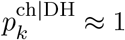 1 if the DH attractor is already deep enough to be self-reproducing under baseline re-challenge.

### 5.2 Likelihood, fit targets and transfer tests

Let 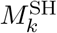 and 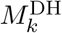 denote the numbers of immediate SH and immediate DH worms forwarded to the challenge assay, and let 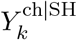 and 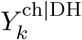 be the corresponding DH counts after re-challenge.

Conditionally on those forwarded counts,

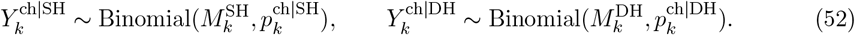

The full likelihood is then

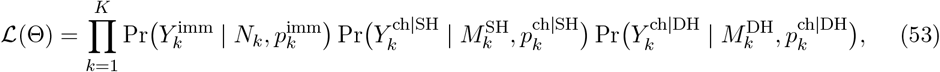

with parameter vector

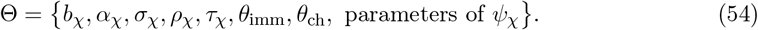

In a first fit one can reduce dimensionality by fixing the latent scale *σ*_χ_ = 1, representing *ψ*_χ_(*t*) by one or two basis functions, or replacing it by free condition-level effects *x*_k_ when only endpoint data are available. If extra-binomial variation is present, the binomial factors in Equations (48) and (52) can be replaced by beta-binomial terms without changing the geometric interpretation.

For the reduced empirical programme used below, it is helpful to separate four roles explicitly:

1. **Primary fit targets:** the 8-OH immediate DH fraction and the 8-OH challenge DH fraction among immediate SH worms;
2. **Consistency check:** the near-absorbing behavior of immediate DH worms under water re-cut, summarised here by setting 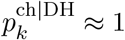 in the reduced model;
3. **Condition-setting data:** the immediate DH fractions under nigericin and monensin, used only to place those treatments on the same latent write axis;
4. **Held-out predictions under the reduced model:** the challenge DH fractions among immediate SH survivors of the nigericin and monensin treatments.

The key point is that the nigericin and monensin *challenge* penetrances do not enter the fit. Only their immediate DH penetrances are used, via the shared immediate threshold, to infer treatment-specific latent means.

A natural staged programme is therefore: fit Equation (53) to the 8-OH challenge data in [4]; then hold the readout geometry fixed and place depolarising perturbations from [5] on the same latent axis using only their immediate DH penetrances. Success in that staged transfer test would support the claim that the reduced rank-one projection is extracting a reusable hidden coordinate rather than merely re-fitting one dataset at a time. We then ask a stricter follow-up question: can a grounded local electrodiffusive feature map recover the same treated-family ordering and similar transfer predictions without reintroducing free per-condition write effects?

Operationally, the reduced fit proceeded in four steps: (i) convert the published rounded percentages into nearest-integer counts; (ii) fix *σ*_χ_ = 1 and *θ*_imm_ = 0 to set the latent scale; (iii) maximize the binomial log-likelihood for (*µ*_8OH_, *µ*_Nig_, *µ*_Mon_, *θ*_ch_); and (iv) report Wald standard errors from the inverse observed Hessian, with approximate 95% probability intervals obtained by the delta method.

**Table 1:** How the reduced calibration was performed. Published rounded percentages were converted to nearest-integer counts at the stated sample sizes. The reduced model fixed the immediate threshold at *θ*_imm_ = 0 and latent variance at 1, then maximized the binomial likelihood in Equation (53) for the remaining reduced parameters.

| Dataset / quantity | Count used | Role | Parameter(s) informed | Model entry |
| --- | --- | --- | --- | --- |
| 8-OH immediate DH | 143 / 573 | primary fit target | $\mu_{8\text{OH}}$ | $p_{8\text{OH}}^{\text{imm}} = \Phi_{\text{N}}(\mu_{8\text{OH}})$ |
| 8-OH challenge DH among immediate SH after water re-cut | 36 / 155 | primary fit target | $\theta_{\text{ch}}$ jointly with $\mu_{8\text{OH}}$ | $p_{8\text{OH}}^{\text{ch\vert SH}}$ from Equation (55) |
| 8-OH challenge DH among immediate DH after water re-cut | 100 / 100 | consistency check | none in reduced fit | used to justify setting $p_k^{\text{ch\vert DH}} = 1$ |
| Nigericin immediate DH | 17 / 132 | condition-setting | $\mu_{\text{Nig}}$ | $p_{\text{Nig}}^{\text{imm}} = \Phi_{\text{N}}(\mu_{\text{Nig}})$ |
| Monensin immediate DH | 11 / 89 | condition-setting | $\mu_{\text{Mon}}$ | $p_{\text{Mon}}^{\text{imm}} = \Phi_{\text{N}}(\mu_{\text{Mon}})$ |
| Nigericin / monensin challenge DH among immediate SH | not used | held-out prediction | none | predicted only after constraining the shared thresholds from 8-OH |

### 5.3 A first reduced fit to published counts

Because the currently available datasets report phenotype counts but not time-resolved estimates of *c*_χ_(*t*) or the kernel *ψ*_χ_(*t*), the first empirical test of the framework should use a deliberately reduced latent-threshold model. All numerical estimates in this section should therefore be interpreted as approximate: the analysis reconstructs counts from rounded percentages reported in the published papers and is intended as a proof-of-concept identifiability test rather than a definitive fit to raw experimental data. The Wald intervals reported below condition on this reconstructed count table and therefore do not propagate the additional uncertainty introduced by reversing rounded percentages.

We collapse each condition to a scalar latent coordinate and fix the latent scale by setting *σ*_χ_ = 1 and *θ*_imm_ = 0. In addition, for this reduced fit we set *ρ*_χ_ = 1 and *τ*_χ_ = 0, so that the challenge assay acts as a lower threshold on the same latent coordinate used for the immediate readout. The resulting model is intentionally minimal.

For condition *k* we write

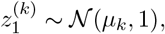

so the immediate double-headed probability is simply

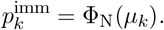

For immediate single-headed worms subjected to a standard water re-cut, we use a lower challenge threshold *θ*_ch_ *<* 0, giving

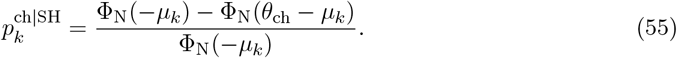

In this reduced fit we treat immediate DH worms as already lying in a deep DH basin and set

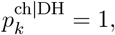

consistent with the 100/100 water re-cut result for immediate DH worms reported by [4].

Using the 8-OH challenge data of [4], and approximating the published rounded percentages by nearest integer counts for the stated sample sizes, we take as calibration targets 143/573 immediate DH after 8-OH and 36/155 challenge DH among immediate SH after water re-cut. Because the 8-OH paper also reports a small residual class of atypical eyeless morphologies, we exclude that ∼ 3% class from the reduced binary fit and treat the calculation as a proof-of-concept rather than a final generative model.

These two 8-OH targets constrain the latent position of the 8-OH treatment and the lower challenge threshold in the reduced model:

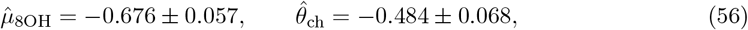

with Wald standard errors. On this normalised latent axis, the immediate DH threshold is at 0 by convention, while the challenge threshold lies about 0.48 latent units below it. In the reduced model, this gap corresponds to a fitted reduced-model *cryptic band* : regenerative states that are subthreshold for immediate visible DH morphology yet suprathreshold for abnormal challenge outcome. The reduced fit reproduces the 8-OH calibration targets,

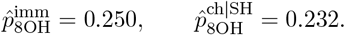

We then use the immediate depolarisation datasets reported for nigericin and monensin by Durant et al. [5] only to locate those treatments on the same latent axis, not to fit any new readout thresholds. The immediate DH penetrances 17/132 for nigericin and 11/89 for monensin imply

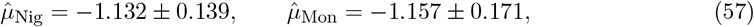

again with approximate Wald standard errors. Substituting these latent means into Equation (55) with the shared threshold from Equation (56) yields the held-out challenge predictions

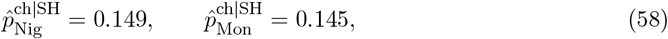

with approximate 95% intervals [0.082, 0.216] and [0.071, 0.218], respectively. These nigericin and monensin re-cut penetrances are *out-of-sample predictions under the reduced model, not fit targets*: no re-challenge data for those treatments enter the calibration. Under this reduced fit, nigericin and monensin are estimated to write smaller shifts than 8-OH along a common latent axis: they generate fewer immediate DH worms, but the model still predicts a non-zero cryptic fraction that should be revealed by standardized re-cutting of the apparently normal survivors.

This reduced fit does not yet identify the full open-path TAS kernel *ψ*_χ_(*t*), the retention factor *ρ*_χ_, or the challenge noise *τ*_χ_. Its role is narrower but important: it shows that published phenotype counts already constrain a nontrivial shared hidden-state geometry consisting of a visible threshold, a lower challenge threshold, and treatment-dependent write amplitudes on a common latent regenerative coordinate. It should therefore be read as a count-based reference layer: a proof-of-concept calibration of a compressed rank-one projection of the full framework, not as empirical validation of the full metric or co-metric structure.

### 5.4 An illustrative grounded in-silico bridge from *G* to phenotype prediction

To show that the biophysical grounding of *G* is tractable rather than merely deferred, we constructed a local in-silico example using a BETSE/BIGR-type electrodiffusive model for one cut geometry and a small family of early-window perturbations [14]. The full construction is given in the Appendix. Briefly, deviations in transmembrane current, a wound-edge gap-junction flux proxy, and Na/K-ATPase activity define a normalized excess-cost proxy in relative units. Finite-difference perturbations then yield explicit local matrices *Q*_χ_, *R*_χ_, and 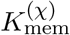. In this illustrative case, the dominant cheap-write direction is concentrated almost entirely in wound-edge gap-junction contrast rather than in cluster-mean *V*_mem_ displacement. Because the current, gap-junction, and pump blocks are normalized by baseline RMS scales, the exact anisotropy is normalization-dependent and should be read as one explicit local instantiation rather than a uniquely identified biological decomposition.

**Table 2:** Reduced fit: parameter estimates, fitted probabilities and held-out predictions. Parameter uncertainties are Wald standard errors from the inverse observed Hessian. Probability intervals are approximate 95% delta-method intervals. Immediate DH means double-headed immediately after the first regeneration round; challenge DH means double-headed after standardized water re-cut of immediate SH survivors.

| Quantity | Estimate | Status | Interpretation |
| --- | --- | --- | --- |
| $\mu_{8OH}$ | $-0.676 \pm 0.057$ | fitted | latent write mean for 8-OH |
| $\theta_{ch}$ | $-0.484 \pm 0.068$ | fitted | challenge threshold; the immediate threshold is fixed at 0 |
| $\mu_{Nig}$ | $-1.132 \pm 0.139$ | condition-setting | latent mean implied by nigericin immediate DH penetrance alone |
| $\mu_{Mon}$ | $-1.157 \pm 0.171$ | condition-setting | latent mean implied by monensin immediate DH penetrance alone |
| $\hat{p}_{8OH}^{imm}$ | 0.250 [0.214, 0.285] | fitted target | reproduces 143/573 immediate DH under 8-OH |
| $\hat{p}_{8OH}^{ch\vert SH}$ | 0.232 [0.166, 0.299] | fitted target | reproduces 36/155 challenge DH among 8-OH immediate SH worms |
| $\hat{p}_{Nig}^{ch\vert SH}$ | 0.149 [0.082, 0.216] | held-out prediction | predicted challenge DH fraction for nigericin immediate SH survivors |
| $\hat{p}_{Mon}^{ch\vert SH}$ | 0.145 [0.071, 0.218] | held-out prediction | predicted challenge DH fraction for monensin immediate SH survivors |

To obtain a semimechanistic phenotype layer, we replaced the free condition-level write effect by a linear readout from a small bank of simulator-derived summaries,

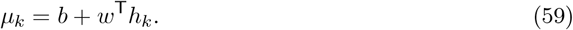

The feature subset was selected by leave-one-treated-out latent prediction error under a soft untreated control anchor 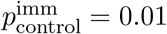. The best stable two-feature subset was

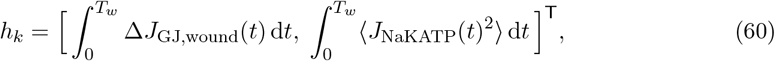

that is, integrated wound-edge gap-junction contrast together with integrated Na/K-ATPase load. Thus the enriched readout combines one state/history feature and one energetic feature.

For this proof-of-concept mapping we associate the experimental families with the closest local perturbation classes in the in-silico model: 8-OH with the local gap-junction block case, nigericin with the local Na-permeability case, and monensin with the local mixed Na/K case. This identification is illustrative rather than unique; its purpose is to test whether a grounded electrodiffusive feature map can recover the same latent ordering and similar held-out challenge scale as the reduced free-*x*_k_ fit.

**Table 3:** Reduced versus illustrative semimechanistic transfer predictions. The semimechanistic bridge uses the enriched simulator-derived readout in Equation (60). The untreated control is included only in the semimechanistic layer, where it is anchored softly at 1% immediate DH for this proof of concept.

| Quantity | Reduced fit | Semimechanistic bridge | Comment |
| --- | --- | --- | --- |
| $\hat{p}_{\text{Nig}}^{\text{ch\vert SH}}$ | 0.149 | 0.148 | similar transfer scale |
| $\hat{p}_{\text{Mon}}^{\text{ch\vert SH}}$ | 0.145 | 0.146 | similar transfer scale |
| $\hat{p}_{\text{control}}^{\text{imm}}$ | not modelled | 0.010 | soft control anchor |

The fitted immediate-layer means are

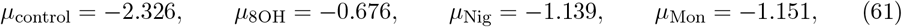

with fitted challenge threshold

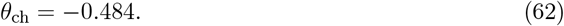

This yields semimechanistic challenge predictions

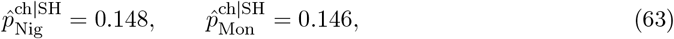

which are close to the reduced free-*x*_k_ predictions in Equation (58). Leave-one-treated-out prediction errors remained below 0.014 latent units, and sweeping the soft control anchor over 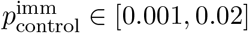 changed the treated-family challenge predictions only at the third decimal place.

This semimechanistic bridge should not be read as independent validation of the reduced fit. Given the small number of treated families, the soft control anchor, and the flexibility of the enriched readout, its role is narrower: it shows that a mechanistically interpretable simulator-derived feature subset can recover the same treated-family ordering and does not contradict the reduced count-based fit. The untreated control anchor is presently a modelling prior rather than a fitted experimental control count, and the perturbation mapping is intentionally local and simplified. The value of this layer is therefore illustrative and methodological: it shows explicitly how a grounded local *G* can be propagated forward through an enriched simulator-derived readout to a phenotype layer without reintroducing free per-condition write amplitudes.

## 6 Predictions from the reduced and semimechanistic latent geometry

The framework yields several experimentally testable predictions. These predictions play different roles. Prediction 6.1 is the near-term falsification test of the reduced rank-one fit. By contrast, Predictions 6.4 and 6.5 are the strongest tests of the specifically geometric content of TAS, because a generic latent-variable account does not by itself imply cut-dependent writable registers or anisotropic rewriting. Prediction 6.2 is sharpened by the grounded in-silico bridge, which provides an explicit local compensatory-cancellation example. The sharpest immediate biological test nevertheless remains the held-out re-challenge of apparently normal survivors from other depolarising interventions.

### 6.1 Prediction 1: held-out re-challenge of nigericin and monensin survivors

#### Protocol

Generate nigericin and monensin cohorts under the same cut geometry used for the immediate DH assays, isolate the apparently normal immediate single-headed survivors, and subject them to the same standardized water re-cut used in the 8-OH cryptic-state experiment.

#### Prediction

Using the shared latent thresholds constrained by the 8-OH calibration and the observed immediate DH penetrances under nigericin and monensin, the reduced model predicts

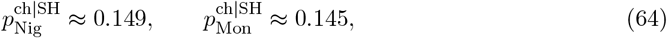

with the intervals reported in Equation (58). The grounded semimechanistic bridge places the same penetrances at

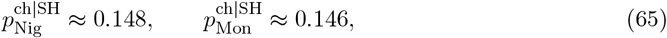

so both empirical layers locate the expected re-challenge penetrance near 15% for each treatment. These re-challenge penetrances are *not* fit targets in either layer: no nigericin or monensin challenge data are used in calibration. Their confirmation or failure therefore provides the cleanest near-term falsification test of the framework.

### 6.2 Prediction 2: matched projected write component, compensatory cancellation, not raw temporal profile

#### Protocol

Compare five perturbation schedules applied during the early sensitive window of the same cut type: (i) constant depolarising exposure, (ii) a short high-amplitude pulse, (iii) a ramped profile, (iv) a sign-reversing protocol in which an early depolarising phase is followed by a matched hyperpolarising counter-phase chosen to cancel the first-order projected write component, and (v) a mild gap-junction co-perturbation chosen to cancel the dominant grounded write coordinate while leaving the bulk depolarising signal approximately intact. Tune the first three protocols so that their first-order projected write component along a dominant local write direction is matched.

#### Prediction

The first three groups should produce the same challenge phenotype to first order, because they write the same hidden displacement *ξ*. Their effective intervention costs should nevertheless differ, with the constant protocol closest to the local efficiency frontier implied by Equation (29). The sign-reversing protocol should behave as a genuine negative control: it expends non-zero effective effort but yields little or no cryptic shift because its leading write component cancels. The grounded in-silico bridge sharpens this logic: the gap-junction compensation group should also suppress hidden writing, even if a bulk depolarisation readout remains comparatively intact, because the dominant cheap-write direction is concentrated in wound-edge gap-junction contrast rather than in mean *V*_mem_ alone.

### 6.3 Prediction 3: a quadratic cost envelope at small amplitude

#### Protocol

Within a single perturbation modality, vary the amplitude over a range that remains below the threshold for gross immediate abnormalities. Measure a quantitative challenge readout, such as the change in DH probability or another standardized phenotype score.

#### Prediction

In the small-amplitude regime, the challenge shift *r*_ch_ should depend approximately linearly on the projected hidden displacement, but the corresponding effective intervention cost must lie above the quadratic envelope

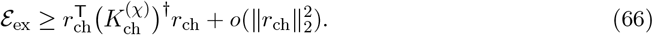

A plot of cost versus challenge-readout magnitude should therefore show a lower envelope with negative controls lying near the origin.

### 6.4 Prediction 4: cut type changes the accessible latent directions

#### Protocol

Apply the same perturbation schedule during regeneration from different cut types, for example pharyngeal, head-only and tail-only amputations. Then re-challenge all groups with a standardized cut and readout.

#### Prediction

Different cut types should induce different local write maps *R*_χ_ and therefore different memory co-metrics 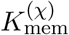. The same perturbation can thus write strongly for one injury geometry and weakly for another, even when the molecular intervention is unchanged. The relevant control parameter is not the amount of tissue exposed to the perturbation, but the geometry of the regenerative episode.

### 6.5 Prediction 5: anisotropic rewriting when dim *Z* ≥ 2

#### Protocol

Use perturbations that preferentially target different aspects of the bioelectric controller, and score challenge outcomes along at least two phenotype axes, for example polarity defects and scaling defects. Combined perturbations can be used to probe mixed directions.

#### Prediction

If the latent controller has dimension greater than one, then 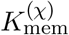 has a non-trivial eigenstructure. Different perturbations will preferentially write into different latent directions, and those directions will be reflected in different phenotype mixtures at matched cost. The experimental signature is not just a stronger or weaker effect, but a rotation of the phenotype mixture consistent with anisotropic access to the hidden state.

### 6.6 Prediction 6: multi-round accumulation and order effects

#### Protocol

Apply a weak sub-threshold perturbation during each of several successive cut-regenerate episodes. In a second version, alternate two distinct weak perturbations in different orders across rounds.

#### Prediction

If retention is high, repeated sub-threshold episodes should accumulate according to Equation (33), eventually crossing the challenge threshold even though no single round is sufficient on its own. If the latent controller is effectively nonlinear, permuting the order of two weak perturbations should yield different challenge outcomes even at matched total dose, providing a biological analogue of non-Abelian transported composition.

## 7 Experimental tests

We outline four concrete experimental protocols suggested by the framework. All are compatible with standard planarian systems and existing bioelectric perturbation methods.

**Experiment 1: immediate held-out transfer test (Prediction 6.1)**. Generate nigericin and monensin cohorts, isolate the immediate SH survivors and subject them to the standardized water re-cut used in the 8-OH cryptic-state assay. Expected outcome: challenge DH penetrances near the values in Equations (58) and (65), that is, roughly 15% for each treatment under both the reduced and semimechanistic layers. Because no such re-challenge data are used in calibration, this is the cleanest immediate falsification test of the framework.

**Experiment 2: temporal profile, compensation, and sign-reversal test (Predictions 6.2– 6.3)**. Use a single cut type and compare five groups during the early sensitive window: constant depolarising exposure, pulsed depolarising exposure, ramped depolarising exposure, a sign-reversing protocol in which a depolarising phase is followed by a matched hyperpolarising counter-phase, and a compensatory protocol in which a mild gap-junction co-perturbation is chosen to cancel the dominant grounded write coordinate while leaving the bulk depolarising signal approximately intact.

Use voltage imaging or an equivalent early marker panel to estimate a common feature vector *f*_k_ and match the projected write component *x*_k_ = *β*^T^*f*_k_ for the first three groups, while verifying near-cancellation for the fourth. Where possible, also measure proxies for wound-edge gap-junction contrast and pump load to test the enriched semimechanistic bridge directly.

Expected outcome: the first three groups produce similar challenge phenotypes but different intervention costs. The sign-reversing group behaves as a negative control despite non-zero effort, and the compensatory gap-junction group also shows strongly reduced challenge shift even if a bulk depolarisation readout remains appreciable.

**Experiment 3: cut-type dependence and anisotropy test (Predictions 6.4–6.5)**. Apply the same perturbation modality across several cut types, and in a parallel series compare several perturbation modalities within one cut type. Score challenge outcomes along at least two phenotype axes, such as polarity and scaling. Expected outcome: challenge penetrance and phenotype mixture vary systematically with cut type and perturbation class, consistent with different 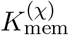 and different dominant write directions.

**Experiment 4: multi-round accumulation test (Prediction 6.6)**. Choose a weak perturbation that is sub-threshold in a single regeneration episode and apply it over *N* = 1, 2, 3, 4, 5 successive cut-regenerate cycles. In a second cohort, alternate two weak perturbations in opposite orders. Expected outcome: the challenge phenotype increases with *N* and, in the order-swap experiment, different sequences produce measurably different outcomes despite matched total exposure.

## 8 Discussion

### 8.1 What the framework adds

The most distinctive contribution of the open-path TAS formulation is directional structure. The cut-dependent co-metric 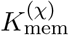 formalises the idea that different injury geometries and perturbation classes need not access the same latent directions with the same efficiency. This turns a qualitative question whether a treatment induces cryptic states into a quantitative one about which latent directions are writable after a given cut. It also yields predictions that are more specifically geometric than the reduced two-threshold fit by itself: cut dependence, anisotropic rewriting, and compensatory protocols that preserve one coarse bioelectric effect while cancelling the dominant hidden write direction.

A second contribution is to separate visible morphology, hidden regenerative state, and challenge readout. In current experimental language these three are often discussed together under the heading of cryptic phenotype. Here they become distinct objects: a coarse anatomical trajectory, a latent endpoint displacement relative to baseline, and a standardized assay that maps that latent displacement to an observable outcome distribution. The reduced count-based fit is the rank-one statistical shadow of this structure, not its entirety.

A third contribution is explicit effective cost accounting. The framework distinguishes the baseline cost of traversing the coarse regeneration trajectory from the excess intervention cost associated with hidden rewriting. In the present version this distinction is no longer purely formal: the local in-silico bridge shows one explicit route by which *G* can be instantiated in relative biophysical units from transmembrane current, gap-junction flux proxies and pump activity, and how the resulting local geometry can be propagated forward to a phenotype layer.

The reduced latent-threshold equations are not claimed to be uniquely TAS-specific. They are the rank-one statistical projection of TAS under current data limitations. The specifically TAS content lies in the geometric definition of hidden state and in the higher-rank predictions of cut dependence, anisotropic rewriting, compensatory cancellation, and order effects.

### 8.2 Relationship to existing models

The framework is complementary to mechanistic bioelectric circuit models of planarian regeneration [12, 14–17]. Those models address the within-episode dynamics of pattern restoration and can predict specific outcomes from specific perturbations. TAS does not replace them. Rather, it provides a higher-level geometry for comparing different physiological realisations of the same coarse regeneration trajectory and for defining the hidden endpoint shifts that challenge experiments reveal.

The new in-silico bridge clarifies how TAS sits on top of BETSE/BIGR-type models. The electrodiffusive model supplies biophysical state variables and local perturbation-response structure; TAS supplies the baseline/excess decomposition, the induced write co-metric, and the challenge-readout layer. What is added is therefore not another simulator of the same kind, but a way to interpret simulator outputs in terms of hidden writing, local efficiency, and transferable readout geometry.

The same is true of information-theoretic viewpoints on biological regulation [21]. Once a challenge readout is treated as a noisy observation of the written hidden state, the cost-information logic developed for mechanical TAS carries over conceptually. What is new in the regenerative setting is that the relevant register is not a kinematic one, but a latent anatomical controller.

### 8.3 Limitations

Five limitations are important. First, the theory assumes that a useful coarse regeneration trajectory *c*_χ_(*t*) can be estimated or approximated from data. Choosing that trajectory is part of the modelling problem, not something given a priori.

Second, the local co-metric results are perturbative. They quantify small hidden displacements around a baseline regenerative episode. Large interventions that cross attractor boundaries are still interpretable in the same language, but the linear formulas become qualitative rather than quantitative.

Third, the latent controller has been treated as finite-dimensional. A realistic planarian pattern-control state is almost certainly distributed in space. The extent to which a low-dimensional reduction captures the relevant biology is an empirical question.

Fourth, the metric *G* is now grounded only locally, in relative units, and through a simulator-based bridge. The present proof-of-concept no longer leaves the grounding problem completely abstract: the main text and Appendix show that a BETSE/BIGR-type model can instantiate *G* and support semimechanistic transfer predictions. But the current bridge still fixes one explicit normalization of current, gap-junction, and pump blocks, so the inferred anisotropy is normalization-dependent; it also uses an illustrative perturbation matching and a soft untreated-control anchor rather than direct physiological calibration.

Fifth, the reduced count-based fit is built from counts reconstructed from rounded percentages. The reported Wald intervals condition on that reconstructed count table and therefore do not propagate the additional uncertainty introduced by reversing rounded percentages.

### 8.4 Outlook

Experimental support for these predictions would establish TAS as more than a descriptive language for regenerative memory. The most immediate quantitative test is the held-out re-challenge of the apparently normal survivors from nigericin and monensin cohorts: under standardised re-cutting, both the reduced count-based layer and the illustrative semimechanistic bridge place the expected challenge DH penetrance near 15%. The predictions that most clearly distinguish TAS from a generic latent-variable account, however, are the cut-dependence and anisotropic-rewriting tests, together with compensatory protocols that preserve a coarse bioelectric effect while suppressing the dominant hidden write direction. In that regime, TAS would provide a quantitative framework for intervention design, allowing timing, amplitude, modality, and sequence to be compared in terms of how efficiently they rewrite the hidden regenerative controller.

More generally, the regeneration application makes explicit a feature of TAS that is only implicit in the mechanical setting. The fundamental geometric object need not be a closed spatial loop; it can instead be an open developmental or physiological episode with a prescribed coarse trajectory, while repetition and challenge convert such episodes into return maps. This suggests natural extensions to other systems with hidden physiological memory, including wound healing, developmental reprogramming, and other forms of anatomical homeostasis. A clear next step is to replace the present soft control anchor, illustrative normalisation choices, and simulator-derived feature weights with direct early-window physiological measurements, thereby linking grounded local geometry more tightly to quantitative phenotype prediction.

## Ethics statement

No new animal experiments were performed for this study.

## Data and code availability

The reduced quantitative analysis uses phenotype counts reconstructed from percentages and sample sizes reported in Durant et al. [4, 5]. All data, code and scripts to reproduce the figures and analyses are available at https://github.com/marcelbtec/tasmorpho

## A Reduced latent-threshold calibration used for the planarian proof-of-principle fit

This section gives the exact statistical model used in the reduced calibration reported in the main text. The aim of this fit is not to estimate the full open-path TAS model developed in the main text, but to test whether a minimal latent-state challenge model can account for the published planarian counts and generate held-out re-challenge predictions. The following two sections then describe two routes beyond this count-only layer: a generic observable proxy based on early physiological measurements, and a concrete local in-silico grounding of *G* in a BETSE-type electrodiffusive model.

**How TAS enters the reduced fit**. The reduced calibration is a lower-dimensional statistical realisation of three specific TAS ingredients rather than a generic latent-threshold model. First, in the full open-path theory a perturbation applied during the prescribed coarse regeneration trajectory writes a hidden endpoint shift

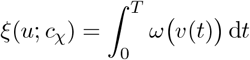

relative to the baseline metric lift. Second, near a fixed baseline episode the local TAS write law has the form

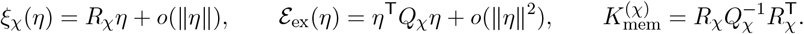

Because the currently available planarian data do not resolve the full coarse trajectory or the full matrix 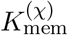, the reduced fit replaces this vector-valued local geometry by a rank-one projection onto a dominant writable coordinate. Writing *ℓ*_χ_ ∈ ℝ^r^ for that effective coordinate, we approximate the hidden shift by the scalar

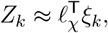

and model its mean as

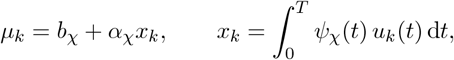

where *x*_k_ is the projected write component of intervention class *k* along an unknown sensitivity kernel *ψ*_χ_(*t*). Third, TAS predicts that immediate morphology and re-challenge are distinct readouts of the same hidden state. In the full theory this appears as two maps,

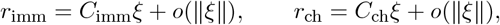

while in the reduced fit these are collapsed to two thresholds on the same scalar latent variable: the immediate threshold at 0 and the lower challenge threshold *θ*_ch_ *<* 0. The fitted interval

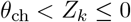

is therefore the scalar analogue of a TAS cryptic channel: invisible to the immediate readout but visible to the challenge readout. This shared hidden-state geometry is what makes held-out transfer possible. Once *θ*_ch_ is fixed from the 8-OH challenge data, nigericin and monensin are not assigned separate challenge parameters; only their locations *µ*_k_ on the common latent write axis change, inferred from their immediate DH penetrances. What the reduced fit does *not* estimate are the full kernel *ψ*_χ_ (*t*), the full co-metric 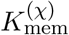, or the continuous-time challenge dynamics. Those objects are collapsed into a shared latent axis and shared thresholds because only endpoint phenotype counts are currently available. The later in-silico section preserves this same threshold layer but replaces the free condition-level write coordinate by simulator-derived features, providing a semimechanistic bridge back to the local TAS objects.

### Data

We used the published planarian counts restated below. For the proof-of-principle fit, when only percentages were reported in the source articles, counts were reconstructed by rounding to the nearest integer for the stated sample sizes. The reduced fit used four aggregated observations:

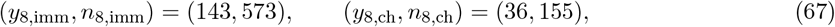

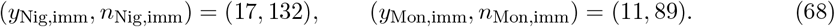

Here “imm” denotes the immediate phenotype after the first regeneration round, “ch” denotes the outcome after the standardized re-amputation challenge, and the challenge count (36, 155) refers to double-headed outcomes among worms that were single-headed in the immediate readout. The reported 100/100 double-headed re-cut outcome for immediately double-headed worms was treated as a consistency check rather than as a fitted data point, since the reduced model assumes that states already beyond the immediate double-headed threshold remain in the double-headed basin under challenge.

### Latent-state model

For each intervention class *k* ∈ {8OH, Nig, Mon}, we introduced a one-dimensional latent regenerative state

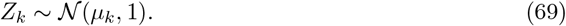

The latent variance was fixed to one to set the scale of the reduced model. The immediate readout threshold was fixed at zero, so that a worm is scored as immediately double-headed if and only if

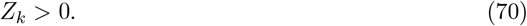

Therefore the immediate double-headed probability under intervention *k* is

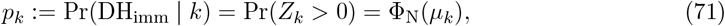

where Φ_N_ is the standard normal cumulative distribution function.

To model cryptic states, we introduced a shared challenge threshold

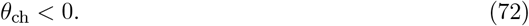

Among worms that are immediately single-headed, i.e. *Z*_k_ ≤ 0, the challenge produces a double-headed outcome when the latent state lies above the challenge threshold:

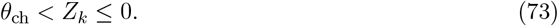

Hence the conditional challenge penetrance is

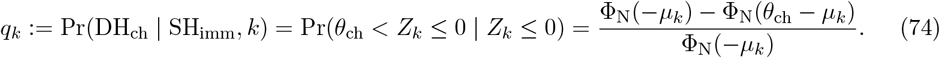

In the fitted dataset, challenge counts were available only for the 8-OH cohort, so *q*_8OH_ was calibrated and *q*_Nig_, *q*_Mon_ were treated as held-out predictions.

### Likelihood

Let

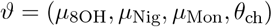

denote the parameter vector. Under the binomial sampling model, the log-likelihood is

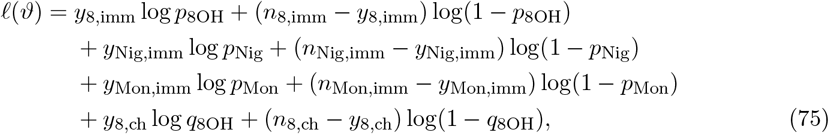

with *p*_k_ and *q*_k_ given by (71) and (74). We estimated 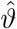 by maximizing (75), equivalently by minimizing the negative log-likelihood. In the numerical implementation this optimization was carried out by Nelder–Mead, although the reduced model also admits the closed-form inversion given below.

### Closed-form structure of the reduced fit

Because each immediate proportion enters through a one-parameter probit map, the reduced fit has a simple inversion structure. The maximum-likelihood estimates of the intervention means are

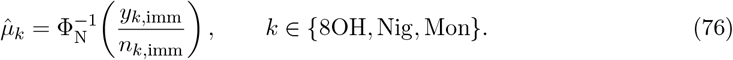

Given 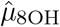, the estimate of the challenge threshold is the unique solution of

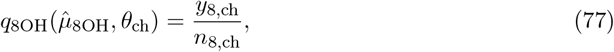

which can be written explicitly as

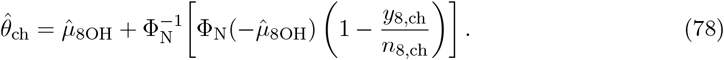

The numerical optimizer reproduced these closed-form values.

### Held-out challenge predictions

Once 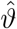 is fixed, the reduced TAS model predicts the re-challenge penetrances for interventions not used in challenge calibration:

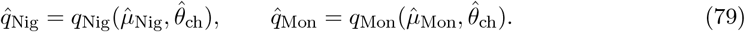

These are the model’s held-out predictions for double-headed outcomes after re-amputating the apparently single-headed survivors of nigericin and monensin treatment.

### Uncertainty quantification

To match the numerical fitting module, uncertainty was estimated from the observed information matrix. Let

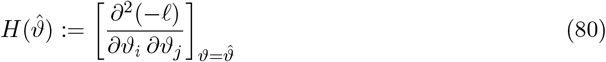

be the Hessian of the negative log-likelihood at the optimum. The covariance matrix was approximated by

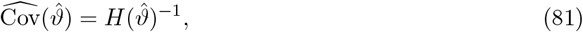

and Wald standard errors were taken as the square roots of the diagonal entries. In the implementation, *H* was computed by finite differences of the negative log-likelihood. If desired, confidence intervals for the held-out probabilities in (79) can then be obtained by the delta method or by parametric bootstrap from 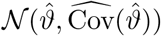

### Interpretation

This reduced calibration should be read as a proof-of-principle bridge between the biological challenge data and the TAS language of hidden state rewriting. It is deliberately lower-dimensional than the full open-path model: the latent coordinate *Z*_k_ summarizes the net hidden shift induced during one regeneration episode, rather than reconstructing the full coarse trajectory *c*_χ_(*t*) or the full write metric. Its main value is to show that published round-1 and round-2 planarian counts already constrain a latent cryptic band and support genuine held-out challenge predictions.

## B Observable and mechanistic proxies for the projected write component

### Generic observable proxy

The reduced fit treats *x*_k_ as a free condition-level effect because only endpoint phenotype counts are currently available. With early-window physiological measurements, one can instead define

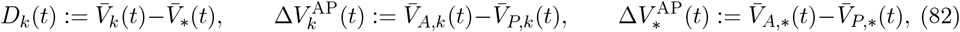

where 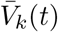 is a chosen whole-fragment voltage statistic under condition *k*, 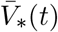 is the corresponding untreated-control statistic, and 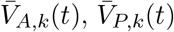 are corresponding anterior and posterior statistics. For an early window [0, *T*_w_], define

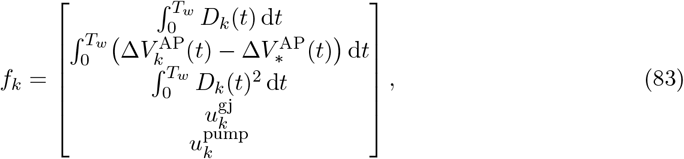

where 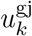 and 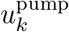 encode gap-junction and pump/ionophore load. One may then write

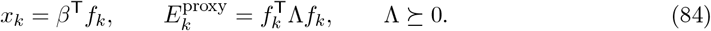

This does not identify the full metric *G*, but it provides a measurable low-rank proxy that can be fit jointly to early bioelectric data and later challenge outcomes. The next section gives one explicit electrodiffusive realization of this logic in a local BETSE example, and then couples the resulting simulator-derived features back to the same latent phenotype layer used in the reduced fit.

### B.1 Local in-silico grounding of *G* and semimechanistic readout bridge

#### BETSE example: a local biophysical instantiation of *G*

To make the metric *G* concrete in relative biophysical units, we ran a reduced BETSE simulation on a single cut geometry using one untreated control and a small family of early-window perturbations. BETSE implements multicellular electrodiffusion with membrane voltage, ion concentrations, gap-junction coupling, and pump dynamics. In the present example, the seeded cluster contained 42 cells before cutting and 39 after the cut event. We used a short early simulation window and compared three perturbation classes: a mild increase in Na^+^ membrane permeability (*e*_Na_), a partial block of gap-junctional coupling (*e*_GJ_), and a hyperpolarizing K^+^ permeability increase used as an auxiliary comparison. Because in this very small local example the Na and K directions were nearly collinear in the raw quadratic estimate, the reported local matrices use the stable two-direction basis (*e*_Na_, *e*_GJ_), while the K^+^ perturbation is retained only as an auxiliary comparison case.

Let *I*_mem_(*t*) denote the vector of BETSE transmembrane current densities, let *J*_NaKATP_(*t*) denote the vector of Na/K-ATPase rates, and define a gap-junction proxy

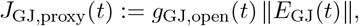

where *g*_GJ,open_ is the relative gap-junction open fraction and *E*_GJ_ is the BETSE gap-junction electric field. Relative to the untreated control, we then defined the normalized excess-cost proxy

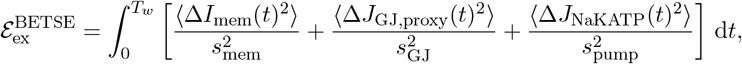

where *s*_mem_, *s*_GJ_, *s*_pump_ are baseline RMS scales and ⟨·⟩ denotes averaging over membrane segments. This defines a local quadratic effort form in relative units.

As a low-dimensional endpoint map for the local write geometry, we used

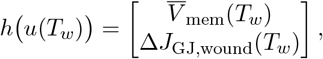

with 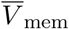 the cluster-mean membrane voltage and Δ*J*_GJ,wound_ the wound-edge minus non-wound gap-junction proxy contrast. Using *e*_Na_ and *e*_GJ_ as the two local perturbation directions, the resulting illustrative matrices were

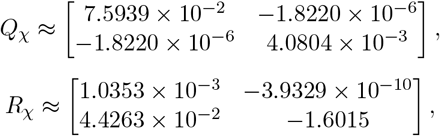

so that

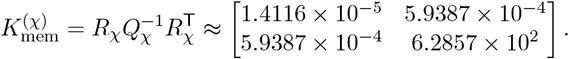

The dominant eigenvector of 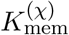 is numerically almost aligned with the wound-edge gap-junction coordinate, indicating that the dominant cheap-write direction in this illustrative local example lies almost entirely in wound-edge gap-junction contrast. This is not a claim that BETSE has identified the true planarian *G*; rather, it shows explicitly how a mechanistic electrodiffusive model can be converted into a concrete local TAS cost matrix and induced memory co-metric in relative biophysical units.

#### A local compensatory prediction from the grounded matrices

For small perturbations *η* = (*η*_Na_, *η*_GJ_)^T^, the hidden endpoint shift is approximated by

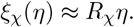

Because the dominant write coordinate is concentrated almost entirely in the second endpoint component, a compensatory protocol can be chosen to preserve the Na-directed bulk depolarization shift while cancelling the dominant hidden write. For a unit Na perturbation, solving

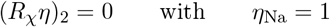

Gives

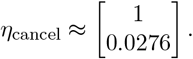

In this local linear approximation,

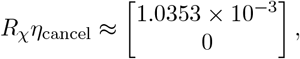

so the wound-edge hidden coordinate is cancelled while the mean-*V*_mem_ shift is essentially unchanged. Conversely, the same dominant hidden write produced by the unit Na perturbation can be matched by a pure gap-junction perturbation

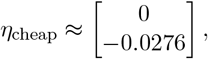

at local cost about 3.1 × 10^−6^, compared with 7.59 × 10^−2^ for the unit Na case. This illustrates the strong anisotropy encoded by 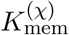 in the local example and provides the mechanistic basis for the compensatory-cancellation prediction stated in the main text.

#### Semimechanistic readout bridge

A semimechanistic extension of the reduced readout can be obtained by replacing the free condition-level write effect by a fitted linear map from a BETSE-derived feature vector,

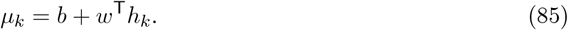

To illustrate the bridge back to the planarian phenotype layer, we associated the local BETSE gap-junction perturbation with the 8-OH condition, the Na-permeability perturbation with the nigericin-like condition, and the mixed Na/K perturbation with the monensin-like condition. This identification is illustrative rather than unique; its purpose is to test whether a grounded electrodiffusive feature map can recover the same latent ordering and similar held-out challenge predictions as the reduced free-*x*_k_ fit.

In a first enriched implementation, *h*_k_ was drawn from a small bank of simulator-derived endpoint and early-window features and the subset was selected by leave-one-treated-out prediction error under a soft near-zero control anchor. The best stable subset was

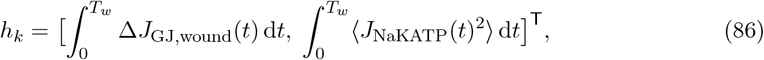

that is, integrated wound-edge gap-junction contrast together with integrated squared Na/K-ATPase load. Using a soft control anchor 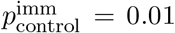, the fitted immediate-layer means were

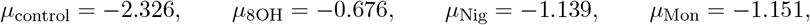

with fitted challenge threshold

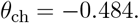

This yields semimechanistic challenge predictions

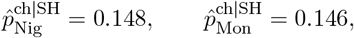

which are very close to the reduced free-*x*_k_ predictions in the main text, while keeping the untreated control near zero in the immediate layer by construction. Leave-one-treated-out errors remained below 0.014 latent units, and sweeping the soft control anchor over 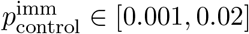 changed the treated-family challenge predictions only at the third decimal place. Because the control anchor is presently a modelling prior rather than a fitted experimental count, because it constrains only the immediate untreated baseline, and because the biological-to-simulator correspondence above is illustrative, this should be read as a semimechanistic proof of concept rather than a definitive phenotype model.

#### Interpretation

Taken together, the two in-silico layers show two distinct things. First, the local BETSE example demonstrates that the TAS metric *G* can be instantiated as a concrete electrodiffusive quadratic form in relative biophysical units and converted into explicit local matrices *Q*_χ_, *R*_χ_, and 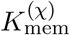. Second, the enriched readout bridge shows that a small simulator-derived feature map can be coupled back to the same latent phenotype layer used in the reduced planarian fit, recovering the treated-family ordering and challenge predictions without assigning free write amplitudes independently to each condition.

## Notes

https://github.com/marcelbtec/tasmorpho

### Competing Interest Statement

The authors have declared no competing interest.

